# Cortical microarchitecture supports preadolescent functional brain network connectivity

**DOI:** 10.64898/2026.09.07.749542

**Authors:** Katherine L. Bottenhorn, Kirthana Sukumaran, Jessica Morrel, Carlos Cardenas-Iniguez, Megan M. Herting

## Abstract

A century ago, Brodmann suggested that local cortical microarchitecture dictates function, yet modern structure– function mapping overwhelmingly focuses on white matter macroarchitecture. Here, we reframe structure-function coupling by linking gray matter microarchitecture and macroscale connectomics, demonstrating how gray matter cellular and neurite density are linked to functional network organization in preadolescents. By applying partial least squares correlation to cortical microarchitecture (i.e., cellular and neurite density) estimates from diffusion MRI and network connectivity estimates from functional MRI, in 6,320 children ages 9-11 years from the U.S.-wide Adolescent Brain Cognitive Development (ABCD) Study, we identified several principles of structure-function coupling. First, we found that functional connectivity is associated with one dominant and global pattern of cellular density differences explaining nearly 80% of shared variance, but by several patterns of neurite density differences. Second, we found similar gradients of sensorimotor network connectivity related to this dominant cellular density pattern and the secondary, rostro-caudal neurite density pattern (explaining 17% of shared variance). Third, we found a profile of attention network connectivity associated with the primary neurite density pattern (44% of shared variance) and the secondary, rostro-caudal cellular density pattern (5% of shared variance). Altogether, this work suggests that cellular density (i.e., neuronal cell bodies, support cells) across cortex largely supports a sensorimotor network connectivity gradient. Conversely, there are several connectivity-related patterns of neurite density (i.e., axons, dendrites) across cortex, providing a glimpse into how local neurite connectivity may differentially support long-range functional connectivity of sensorimotor networks and attention networks.

## Introduction

Early twentieth century researchers mapped the human brain’s cyto-and myeloarchitecture, assuming regional differences reflected specialized function (Brodmann, 1909; Campbell, 1905; Economo & Koskinas, 1925; Foerster, 1934; Smith, 1907; Vogt & Vogt, 1919). One feature of this architecture is neuronal density, which follows a rostro-caudal gradient in mammalian cortex, with higher density in caudal regions like the occipital lobe and lower density in rostral regions like the prefrontal cortex (Hilgetag et al., 2019; Huntenburg et al., 2018). Some speculate that this represents a computational gradient, ranging from many processing units in primary visual and somatomotor cortex to fewer, but more densely connected, units in higher-order cortex (Cahalane et al., 2012; Charvet & Finlay, 2014; Hilgetag et al., 2019). With the advent of magnetic resonance imaging (MRI), cognition and behavior have been linked to brain macro-and cytoarchitecture, often leveraging cytoarchitectonic labels commonly used to localize brain activation in functional task-based MRI (fMRI) studies (Amunts et al., 2007; Devlin & Poldrack, 2007; Triarhou, 2007; Van Essen & Dierker, 2007). Functional MRI data has also been used to assess functional variability within architectonic areas (Ardila et al., 2015; Bludau et al., 2014; Hampson et al., 2006; Zilles & Amunts, 2010). As the field has shifted focus from localizing function to individual brain regions towards a network-based view, wherein functions emerge from interactions among sets of brain regions (Pessoa, 2014), fMRI has been used to identify distributed functional brain networks (van den Heuvel & Hulshoff Pol, 2010) that are consistent across individuals, states of consciousness, and tasks (S. M. Smith et al., 2009). These networks correspond to distinct cognitive processes and behaviors (Laird et al., 2011) and are supported in part by white matter tracts that can be measured with diffusion-weighted MRI (dMRI) (Sui et al., 2014).

Multimodal MRI data can link brain structure and function *in vivo.* Much of this work has focused on examining white matter connections between spatially distinct regions of large-scale functional networks, linking structural and functional *connectivity* (SC and FC, respectively) (Honey et al., 2009; Liégeois et al., 2020). Using graph theory approaches, researchers have uncovered shared topological properties between structural and functional brain networks, such as small-worldness and rich clubs organizations. They have found stronger structure-function correspondence in visual, motor, and default mode networks than in subcortical, limbic, and attention networks (Bassett et al., 2006; Grayson et al., 2014; Greicius et al., 2009; Heuvel et al., 2008; Horn et al., 2014; Osmanlıoğlu et al., 2019; Sporns & Zwi, 2004; Wang et al., 2015). SC-FC correspondence also appears to follow a functional gradient, from unimodal sensory and motor areas to transmodal association areas, with weaker structure-function coupling in the transmodal cortex (Baum et al., 2020). This coupling further changes over development, and age-related differences in both structural and functional MRI measures tend to follow the same unimodal-to-transmodal axis (Bottenhorn et al., 2025; Sydnor et al., 2021, 2023). Consistent with this, longitudinal work suggests sensory and motor cortex mature earlier than association cortex (Gogtay et al., 2004).

Cortical development during childhood and adolescence involves ongoing synaptic pruning, myelination, and apoptosis (Huttenlocher & Dabholkar, 1997), alongside the continued maturation of large-scale functional networks (Grayson & Fair, 2017). Multimodal neuroimaging studies suggest that how brain structure and function relate to one another changes across this period, and that these changes differ along a unimodal-to-transmodal neurofunctional axis (Baum et al., 2020; Park et al., 2022; Soman et al., 2023). Specifically, structure-function correspondence is stronger in sensory and motor regions early in childhood and weakens with age, while structural and functional connectivity in association cortex increase with age (Baum et al., 2020; Park et al., 2022). Additionally, structural differentiation between brain regions (e.g., cortical microarchitecture similarity, white matter connectivity) follows the same unimodal-to-transmodal pattern in childhood, and age-related increases in in large-scale network differentiation are accompanied by weaker functional connectivity between networks (Park et al., 2022). Altogether, this work suggests a coupling between structural brain differences and the development of functional organization. However, it largely overlooks how differences in local microarchitecture, which reflect ongoing neurobiological processes like synaptic pruning, are related to network-level brain function during development.

This work builds on Brodmann’s assertion that “the specific histological differentiation of cortical areas provides irrefutable proof of their specific functional differentiation” (Brodmann & Garey, 1994), combining more recent evidence that cytoarchitectonic similarity is greater in structurally connected cortical regions (Wei et al., 2018) and the current network model of human brain function (Behrens & Sporns, 2012; Mišić & Sporns, 2016). Using a cross-sectional sample of 6,007 youth enrolled in the Adolescent Brain Cognitive Development℠ Study (ABCD Study®), we identified latent associations between cortical microarchitecture and functional connectivity in large-scale brain networks at ages 9-11 years using partial least square correlation (PLSC) analyses (**Figure 1**). Cortical microarchitecture was estimated using restriction spectrum imaging (RSI), a biophysical model applied to dMRI data that separately resolves intracellular and extracellular compartments (White et al., 2012). Within the intracellular compartment, isotropic diffusion reflects the density of cell bodies (i.e., neurons and glia), while directional diffusion reflects the density of neurites (i.e., axons and dendrites) in cortex. Between the ages of 9 to 13 years, cellular density increases, while neurite density decreases (Bottenhorn et al., 2023) though regional trajectories differ between sensorimotor and associative cortex (Bottenhorn et al., 2025) and functional connectivity within and between large-scale networks also changes over this period (Betzel et al., 2014; Edde et al., 2021; Sun et al., 2025). Mapping cortical microarchitecture to large-scale network topology will help uncover fundamental principles of human brain function, enabling future investigations to study individual differences in connectivity-related microarchitecture and their implications for the maturation of behavior and cognition.

**Figure 1.**
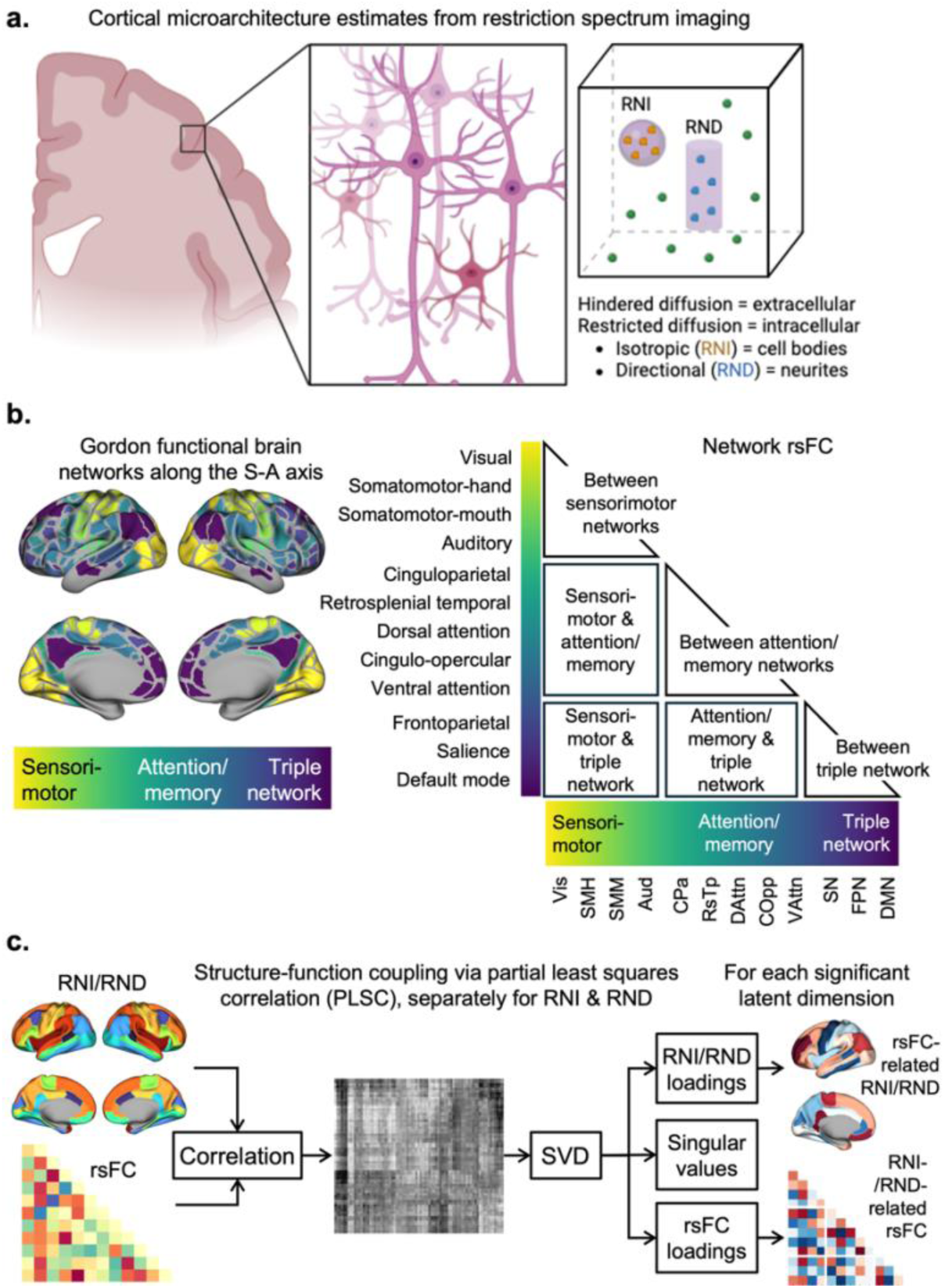
Schematic representation of neural phenotypes and analysis pipeline used to identify modes of structure-function coupling. a) Schematic representation of the two microarchitecture measures used in these analyses (left). For intracellular diffusion, restricted normalized isotropic (RNI) diffusion is thought to represent the density of cell bodies (e.g., neurons and glia), while restricted normalized directional (RND) diffusion is thought to represent the density of neurites (e.g., both axons and dendrites) in gray matter. b) Resting-state functional connectivity (rsFC) is estimated here at the network level, as delineated by Gordon et al. (2016; left). For ease of interpretation, networks were aligned with the sensorimotor-association axis (S-A axis; Sydnor et al., 2021) and an annotated schematic adjacency matrix is displayed (middle). Network abbreviations: visual, Vis; auditory, Aud; somatomotor-hand, SMH; somatomotor-mouth, SMM; dorsal attention, DAttn; ventral attention, VAttn; retrosplenial temporal, RsTp; cinguloparietal, CPa; cingulo-opercular, COpp; salience, SN; frontoparietal, FPN; default mode, DMN. c) Coupling between cortical microarchitecture and functional connectivity was assessed using partial least squares correlation (PLSC), separately for RNI and RND to identify patterns of cellular density and neurite density linked to individual differences in large-scale resting-state functional network connectivity. Prior to PLSC, participant age, sex, handedness, study site, and head motion during MRI scans were regressed out of RNI/RND and rsFC estimates. Briefly, PLSC proceeds by calculating correlations between each pair of variables in the two blocks (here: RNI/RND and rsFC), then performs a singular value decomposition (SVD) on these correlations to identify variable loadings in each block on a set of latent dimensions, as well as the singular values for each dimension. Permutation testing identifies significant latent dimensions (not shown) and bootstrap resampling identifies brain regions exhibiting significantly rsFC-related RNI/RND and brain networks exhibiting significantly RNI-/RND-related rsFC. For more details, see the Methods.

## Methods

### Participants

Cross-sectional data were accessed from the Adolescent Brain Cognitive Development (ABCD) Study’s 5.1 data release (https://dx.doi.org/10.15154/z563-zd24). The ABCD Study enrolled 11,876 children 9 to 10 years of age (mean age = 9.49; 48% female) in a 10-year longitudinal study (Volkow et al., 2018). Participants were recruited at 21 study sites across the United States from elementary schools (private, public, and charter schools) and birth-registries (for twins) in a sampling design that aimed to represent the nationwide sociodemographic diversity (Garavan et al., 2018). All experimental and consent procedures were approved by a central institutional review board and human research protections programs at the University of California San Diego. Each participant provided written assent to participate in the study and their legal guardian provided written agreement to participate. Here, we use a cross-sectional subset of ABCD Study data collected at 9-10 years of age, including magnetic resonance imaging (MRI), in addition to measures of participants’ sex at birth, handedness, and age. Exclusion criteria for the ABCD study included lack of English proficiency, severe sensory, neurological, medical or intellectual limitations, and inability to complete an MRI scan. For this study, we first excluded individuals from site 15 due to hardware issues (*N* = 458); then further excluded participants who had incidental neurological findings from their MRI scans, failed the ABCD Study imaging quality control procedures, or had greater than 2mm of head motion during dMRI scans (*N* = 2,452); and finally excluded participants who had greater than 0.5mm head motion during rs-fMRI scans (*N* = 2,633) for a final sample of 6,320 participants with sufficient quality dMRI and rs-fMRI data and complete data on all covariates. Numbers of participants included and excluded at each step are presented in Supplementary Figure 1, while final sample characteristics for the current study are described in Table 1.

**Table 1.**
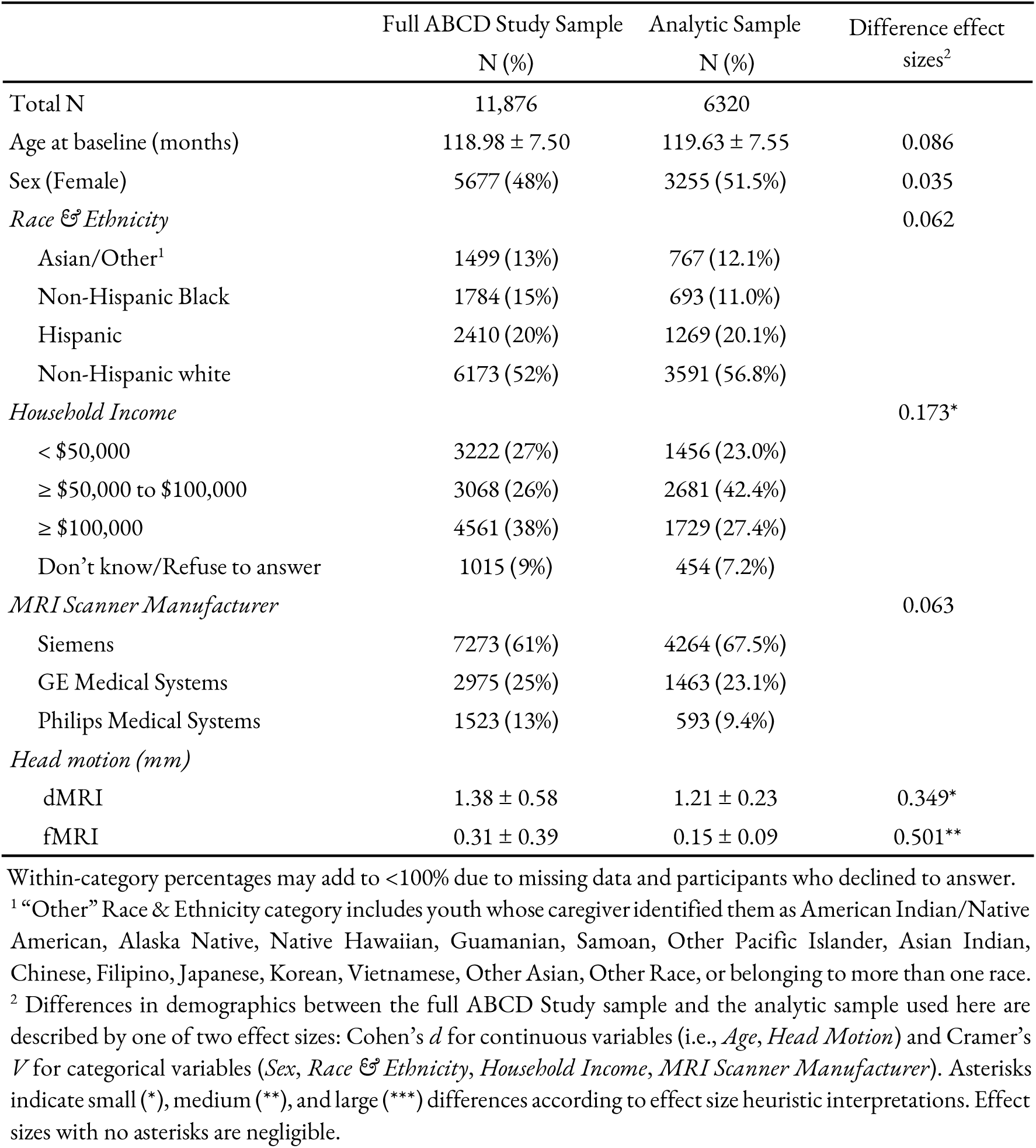
Sample Demographics.

|  | Full ABCD Study Sample<br>N (%) | Analytic Sample<br>N (%) | Difference effect<br>sizes<sup>2</sup> |
| --- | --- | --- | --- |
| Total N | 11,876 | 6320 |  |
| Age at baseline (months) | 118.98 ± 7.50 | 119.63 ± 7.55 | 0.086 |
| Sex (Female) | 5677 (48%) | 3255 (51.5%) | 0.035 |
| *Race & Ethnicity* |  |  | 0.062 |
| Asian/Other<sup>1</sup> | 1499 (13%) | 767 (12.1%) |  |
| Non-Hispanic Black | 1784 (15%) | 693 (11.0%) |  |
| Hispanic | 2410 (20%) | 1269 (20.1%) |  |
| Non-Hispanic white | 6173 (52%) | 3591 (56.8%) |  |
| *Household Income* |  |  | 0.173* |
| < \$50,000 | 3222 (27%) | 1456 (23.0%) | |
| ≥ \$50,000 to \$100,000 | 3068 (26%) | 2681 (42.4%) | |
| ≥ \$100,000 | 4561 (38%) | 1729 (27.4%) | |
| Don't know/Refuse to answer | 1015 (9%) | 454 (7.2%) |  |
| *MRI Scanner Manufacturer* |  |  | 0.063 |
| Siemens | 7273 (61%) | 4264 (67.5%) |  |
| GE Medical Systems | 2975 (25%) | 1463 (23.1%) |  |
| Philips Medical Systems | 1523 (13%) | 593 (9.4%) |  |
| *Head motion (mm)* |  |  |  |
| dMRI | 1.38 ± 0.58 | 1.21 ± 0.23 | 0.349* |
| fMRI | 0.31 ± 0.39 | 0.15 ± 0.09 | 0.501** |

Within-category percentages may add to <100% due to missing data and participants who declined to answer.
<sup>1</sup> "Other" Race & Ethnicity category includes youth whose caregiver identified them as American Indian/Native American, Alaska Native, Native Hawaiian, Guamanian, Samoan, Other Pacific Islander, Asian Indian, Chinese, Filipino, Japanese, Korean, Vietnamese, Other Asian, Other Race, or belonging to more than one race.
<sup>2</sup> Differences in demographics between the full ABCD Study sample and the analytic sample used here are described by one of two effect sizes: Cohen's $d$ for continuous variables (i.e., *Age*, *Head Motion*) and Cramer's $V$ for categorical variables (*Sex*, *Race & Ethnicity*, *Household Income*, *MRI Scanner Manufacturer*). Asterisks indicate small (\*), medium (\*\*), and large (\*\*\*) differences according to effect size heuristic interpretations. Effect sizes with no asterisks are negligible.

### Neuroimaging Data

A harmonized data protocol was utilized across sites with either a Siemens, Phillips, or GE 3T MRI scanner. Motion compliance training, as well as real-time, prospective motion correction was used to reduce motion distortion (B. J. Casey et al., 2018).

#### Diffusion Weighted Imaging (DWI): Acquisition, Processing, and Quality Control

Diffusion-weighted MRI acquisition included a voxel size of 1.7 mm isotropic and implements multiband EPI (Moeller et al., 2010; Setsompop et al., 2012) with slice acceleration factor 3. Each DWI acquisition included a fieldmap scan for B0 distortion correction. The ABCD Study employs a multi-shell diffusion acquisition protocol that includes 7 b=0 frames as well as 96 total diffusion directions at 4 b-values (6 with b = 500 s/mm^2^, 15 with b = 1000 s/mm^2^, 15 with b = 2000 s/mm^2^, and 60 with b = 3000 s/mm^2^). All images underwent distortion correction, bias field correction, motion correction, and manual and automated quality control per the steps detailed by Hagler and colleagues (2019). Only images without clinically significant incidental findings (*mvif_scove* = 1 or 2) that passed all ABCD quality-control parameters were included in analysis (*imgincl_dmvi_include* = 1).

#### DWI Modeling: Restricted Spectrum Imaging (RSI)

Restricted spectrum imaging (RSI) is a more advanced modeling technique that utilizes all 96 directions collected as part of ABCD’s multi-shell acquisition protocol (B. J. Casey et al., 2018; Hagler et al., 2019; White et al., 2012). RSI provides detailed information regarding both extracellular (i.e., from hindered diffusion; Figure 1A) and intracellular (i.e., from restricted diffusion; Figure 1A) compartments within the brain (White et al., 2012, 2014). RSI model outputs include normalized measures, all of which are unitless on a scale of 0 to 1. The current study focuses on two intracellular diffusion metrics of interest: restricted normalized isotropic diffusion (RNI), which is thought to represent the density of cell bodies (e.g., neurons and glia), and restricted normalized directional diffusion (RND), thought to represent the density of neurites (e.g., both axons and dendrites) in gray matter (Palmer et al., 2022; White et al., 2012). Mean RNI and RND were calculated for cortical gray matter ROIs from the Desikan-Killiany atlas (Desikan et al., 2006).

#### Resting State fMRI (rs-fMRI): Acquisition, Processing, and Quality Control

Twenty cumulative minutes of resting-state functional MRI data was collected across two sets of two five-minute acquisition periods, while subjects were instructed to keep their eyes open and fixed on a crosshair. This increases the probability of collecting enough data with low motion per the ABCD Study’s standards (>12.5 minutes of data with framewise displacement (FD) < 0.2 mm) (Power et al. 2014). Resting state scans were acquired using an echo-planar imaging sequence in the axial plane, with the following parameters: TR = 800 ms, TE = 30 ms, flip angle = 90°, voxel size = 2.4 mm^3^, 60 slices. Only images without clinically significant incidental findings (*mvif_scove* = 1 or 2) that passed all ABCD quality-control parameters were included in analysis (*imgincl_vsfmvi_include* = 1). Image processing steps have been previously described by Hagler and colleagues (2019).

#### rs-fMRI: Gordon Parcellation and Functional Connectivity Analysis

Twelve resting-state networks were functionally defined using resting-state functional connectivity, including sensorimotor networks (i.e., visual, auditory, somatosensory-hand, somatosensory-mouth) (Gordon et al., 2016). Intra-network correlations were calculated by averaging the pairwise correlations for ROIs belonging to that network; inter-network correlations were calculated by averaging the pairwise correlations between ROIs within the first network and ROIs within the second network; subcortical-network correlations were calculated by averaging the pairwise correlations between ROIs within a network and a given subcortical ROI (Hagler et al., 2019).

In interpreting networks, they were grouped based on their function into Sensorimotor, Attention/Memory, and Triple Network, which also correspond with their order along the sensorimotor-association axis (S-A, Figure 1B) (Sydnor et al., 2021). Sensorimotor networks represent the sensorimotor pole of the S-A axis and include primary visual (Vis) and auditory (Aud) networks, as well as somatomotor-hand (SMH) and -mouth (SMM) networks. Attention/memory networks fall along the middle of the S-A axis and include cinguloparietal (CPa), retrosplenial temporal (RsTp), dorsal attention (DAN), ventral attention (VAN), and cingulo-opercular (COpp) networks. The triple network model of higher-order functional networks with transdiagnostic links to psychopathology represents the association pole of the S-A axis and includes the salience (SN), frontoparietal (FPN), and default mode (DMN) networks (Menon, 2011).

### Covariates

Covariates regressed out of resting-state functional connectivity (rsFC) and cortical microarchitecture measures prior to partial least squares correlation (PLSC) analysis consisted of precision variables related to both the child and MRI collection, including child’s age during their visit, child’s sex at birth (male or female), handedness (right, left, or mixed), data collection site, scanner manufacturer (Siemens, Philips, GE), and head motion (i.e., average framewise displacement, in mm) to account for scanner differences (i.e., hardware and software) and address EPI sequences’ sensitivity to head motion (Ciric et al., 2018; Power et al., 2012; Satterthwaite et al., 2012).

### Analyses

Descriptive statistics and categorical counts for sociodemographic variables (i.e., age, sex, race/ethnicity, household income), scanner manufacturer, and head motion were calculated in R for both the full ABCD Study sample and the analytic sample analyzed here (Table 1).

Partial Least Square Correlation (PLSC; Figure 1C) was performed to identify shared variance between cortical microarchitecture and functional connectivity using the ‘TExposition’ and ‘TInPosition’ packages in R (version 4.2.1). To do so, this multivariate method PLSC takes two data blocks-in this case, cortical microarchitecture measures (RNI or RND) and functional connectivity measures, over the same set of participants and identifies latent dimensions that maximize the covariance between the two (Krishnan et al., 2011; McIntosh & Lobaugh, 2004). The cortical microarchitecture block included 68 cortical brain regions (i.e., measured as isotropic (RNI) or anisotropic (RND) diffusion) obtained via RSI whereas the functional connectivity block included 78 connections of 12 networks.

Prior to running PLSC, both microarchitecture and rsFC measures were residualized, mean-centered, and normalized (to standard deviation of 1), to remove confounding influences. PLSC breaks down the correlation between the two data blocks into latent dimensions where each latent dimension represents a weighted combination of original variables from both data blocks, capturing shared variance between them. The contribution of each variable to these latent dimensions is represented by latent variable loadings/saliences while the strength of the relationship between the blocks along each latent dimension is captured by singular values and the latent scores represents the participant contributions from each block towards the latent dimension (all derived from a matrix decomposition tool called Singular Value Decomposition (SVD) (Abdi, n.d.). To identify the number of significant latent dimensions a permutation test was performed using Perm4PLSC() function randomly sampling data with 1000 iterations (Abdi & Williams, 2013). The robustness of the identified regional associations for each significant latent dimension were quantified via bootstrapping using the Boot4PLSC() function with 10,000 resamples in package ‘data4PCCAR’’ (Hachay, 2018/2026). As bootstrap ratios are analogous to *z*-scores, any region with a bootstrap ratio greater than 2.5 or less than -2.5 was considered “significantly” contributing to the associated dimension (i.e., *p* < 0.01).

## Results

The analytic sample used here to assess latent coupling between cortical microarchitecture and functional network connectivity was *N* = 6,320 youth (Table 1). Effect sizes comparing the current analytic sample with the full ABCD Study sample are presented in Table 1 (“*Diffevence effect sizes*”). The current analytic sample exhibited negligible differences in age at enrollment, sex at birth, race/ethnicity, and MRI scanner manufacturer from the full ABCD Study sample, small differences in household income and head motion during dMRI scans, and medium differences in head motion during fMRI scans.

Partial least squares correlation identified 10 significant latent dimensions of cellular density-functional connectivity coupling (RNI-rsFC) and 9 significant dimensions of neurite density-functional connectivity (RND-rsFC) coupling (all *p* < 0.01; Table 2). Given this, we focus our results and interpretation on the primary and secondary latent dimensions for both RNI-rsFC and RND-rsFC coupling. The remaining significant dimensions reported in Supplementary Figures 4-16. For each latent dimension, loadings for each brain region and functional connection are presented below as bootstrap ratios, which are analogous to *z*-scores. Directionality of these ratios is relative within each latent dimension, such that brain regions and connections with the same sign (i.e., +/+, -/-) are positively coupled, while brain regions and connections with the opposite sign (i.e., +/-, -/+) are negatively coupled.

**Table 2.** Proportions of shared microstructure-function variance explained by each latent dimension from PLSC, separately for RNI and RND.

| Dimension | RNI-rsFC | RND-rsFC |
| --- | --- | --- |
| 1 | 79% | 44% |
| 2 | 5% | 17% |
| 3 | 3% | 6% |
| 4 | 3% | 6% |
| 5 | 2% | 4% |
| 6 | 1% | 4% |
| 7 | 1% | 3% |
| 8 | 1% | 3% |
| 9 | 1% | 2% |
| 10 | 1% | – |

*Note.* Proportions of variance explained are only shown for significant latent dimensions. Abbreviations: PLSC, partial least squares correlation; RNI, restricted normalized isotropic diffusion (i.e., cellular density); RND, restricted normalized directional diffusion (i.e., neurite density); rsFC, resting state functional connectivity.

### Correspondence between cellular density and functional network connectivity

The primary RNI-rsFC latent dimension explained 79% of the shared variance and was characterized by greater RNI across the cortex underlying a gradient of sensorimotor rsFC (Figure 2a). The highest RNI was in posterior regions and lowest RNI in auditory, cingulate, and ventromedial prefrontal cortex (Figure 2a, left). The sensorimotor rsFC gradient (Figure 2a, middle) was characterized by greater rsFC between sensory and motor networks (e.g., auditory, primary somatosensory), but weaker rsFC between sensorimotor networks and the triple network model (e.g., frontoparietal, salience, default). Attention and memory-related networks’ rsFC with sensorimotor networks was mixed. Specifically, we found RNI-related rsFC was weaker in the cinguloparietal network (i.e., with somatomotor and auditory networks) and cingulo-opercular network (i.e., with somatomotor-hand and auditory networks), but stronger in the ventral attention network (i.e., with visual and somatomotor-hand networks). Similarly, RNI-related within-network rsFC (Figure 2a, right) was stronger in sensorimotor networks and the cinguloparietal network, but weaker in retrosplenial, cingulo-opercular, and ventral attention networks.

**Figure 2.**
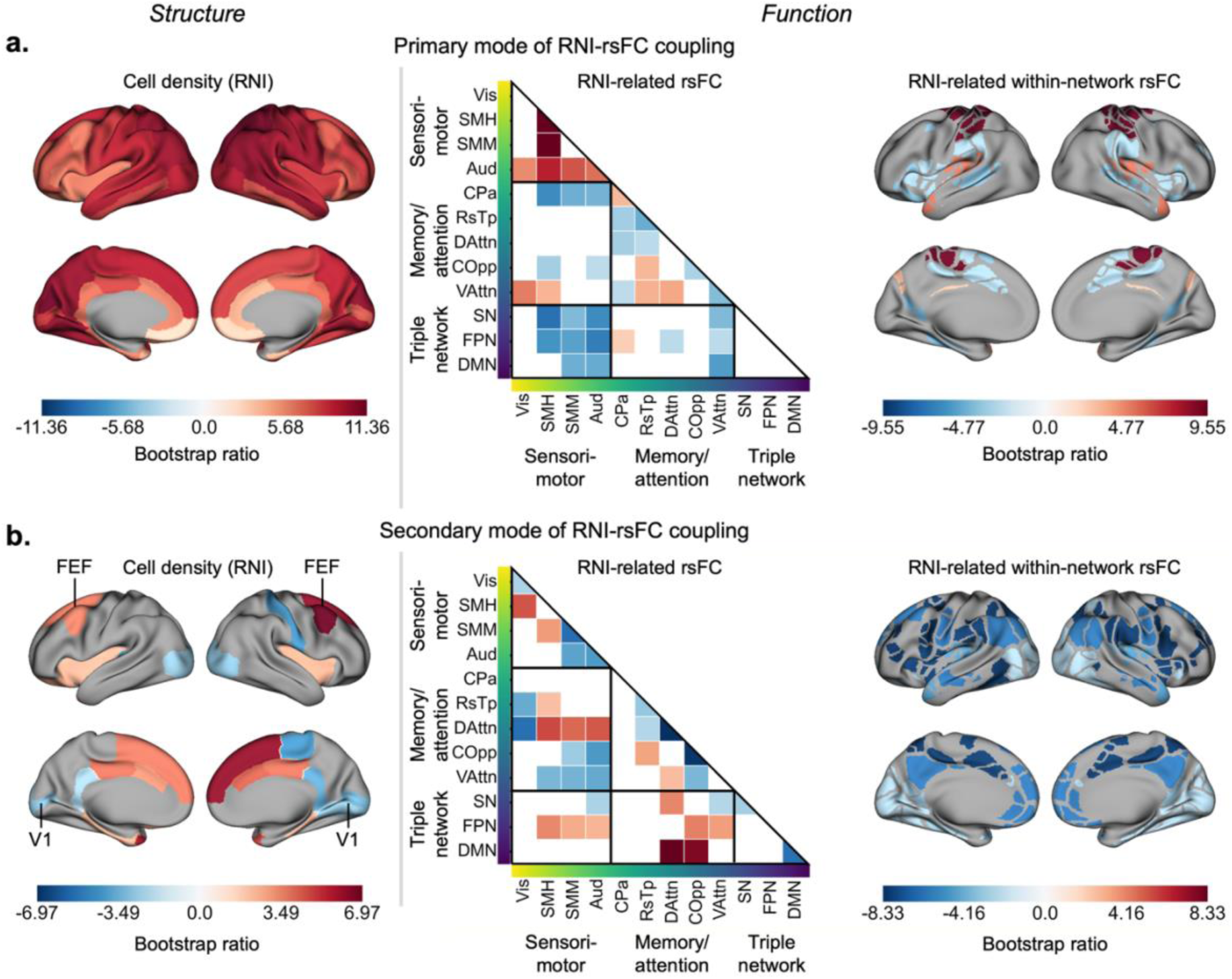
Primary and secondary patterns of coupling between cellular density and functional connectivity. Cellular density patterns are shown on the left, as bootstrap ratios representing brain regions that significantly contribute to the primary (a) and secondary (b) modes of RNI-rsFC coupling. Related patterns of functional connectivity are shown in the middle (all rsFC) and right (within-network rsFC) as bootstrap ratios representing network connections that significantly contribute to the primary and secondary modes of RNI-rsFC coupling. Networks are grouped based on their function into Sensorimotor, Attention/Memory, and Triple Network, which are ordered from unimodal to transmodal along the x-and y-axes in the middle column. A) The primary mode of RNI-rsFC coupling accounted for 79% of the shared variance between cortical RNI and functional network connectivity. A global pattern of higher cellular density (i.e., RNI) was related to a gradient of sensorimotornetwork connectivity that was stronger among sensory and motor networks (Vis, SMH, SMM, Aud), weaker between sensorimotornetworks and the triple network model (SN, FPN, DMN), and mixed between sensorimotornetworks and attention/memory networks (CPa, COpp, VAttn). B) The secondary mode of RNI-rsFC coupling accounted for 6% of the shared variance between cortical RNI and functional network connectivity. A posterior-to-anterior pattern of lower cellular density in posterior regions (i.e., pericalcarine gyrus, V1; lateral occipital, posterior cingulate, postcentral gyrus) and higher cellular density in anterior regions (i.e., insula; midcingulate, superior frontal, caudal middle frontal gyri including frontal eye fields, FEF) was related to largely weaker within-network connectivity (right) and mixed stronger/weaker connectivity in frontoparietal, dorsal attention, and ventral attention networks (middle). Abbreviations: visual, Vis; auditory, Aud; somatomotor-hand, SMH; somatomotor-mouth, SMM; dorsal attention, DAttn; ventral attention, VAttn; retrosplenial temporal, RsTp; cinguloparietal, CPa; cingulo-opercular, COpp; salience, SN; frontoparietal, FPN; default mode, DMN.

The secondary latent dimension of RNI-rsFC coupling explained 5% of shared variance and reflected a pattern of higher-order development (Figure 2b). This pattern included greater RNI in bilateral medial superior frontal, mid-cingulate, insula, and caudal middle frontal gyri, but less RNI in bilateral occipital regions and auditory cortex, as well as right postcentral, paracentral, and posterior cingulate gyri (Figure 2b, left). These RNI differences were accompanied by between-network rsFC (Figure 2b, middle) that was stronger in dorsal attention (except with visual network) and frontoparietal networks (with cinguloparietal, ventral attention, auditory, and somatosensory networks), but weaker in the ventral attention and cingulo-opercular networks. Further, RNI-related within-network rsFC was weaker overall (Figure 2b, right), specifically in auditory, somatosensory, dorsal attention, retrosplenial temporal, cingulo-opercular, salience, and default mode networks.

### Correspondence between neurite density and functional network connectivity

The primary latent dimension of RND-rsFC coupling explained 44% of the shared variance and reflected a widespread pattern of greater RND juxtaposed with sparser rsFC (Figure 3a). This pattern included greater RND across most of the cortex, except for insula, cingulate, calcarine, and superior temporal gyri (Figure 3a, left). This greater RND was accompanied by between-network rsFC (Figure 3a, middle) that was stronger for the dorsal attention and frontoparietal networks, but weaker for the cingulo-opercular and ventral attention networks. RND-related within-network rsFC (Figure 3a, right) was weaker overall (Figure 3a, right), notably in auditory, somatosensory, dorsal attention, retrosplenial temporal, cingulo-opercular, salience, and default mode networks.

**Figure 3.**
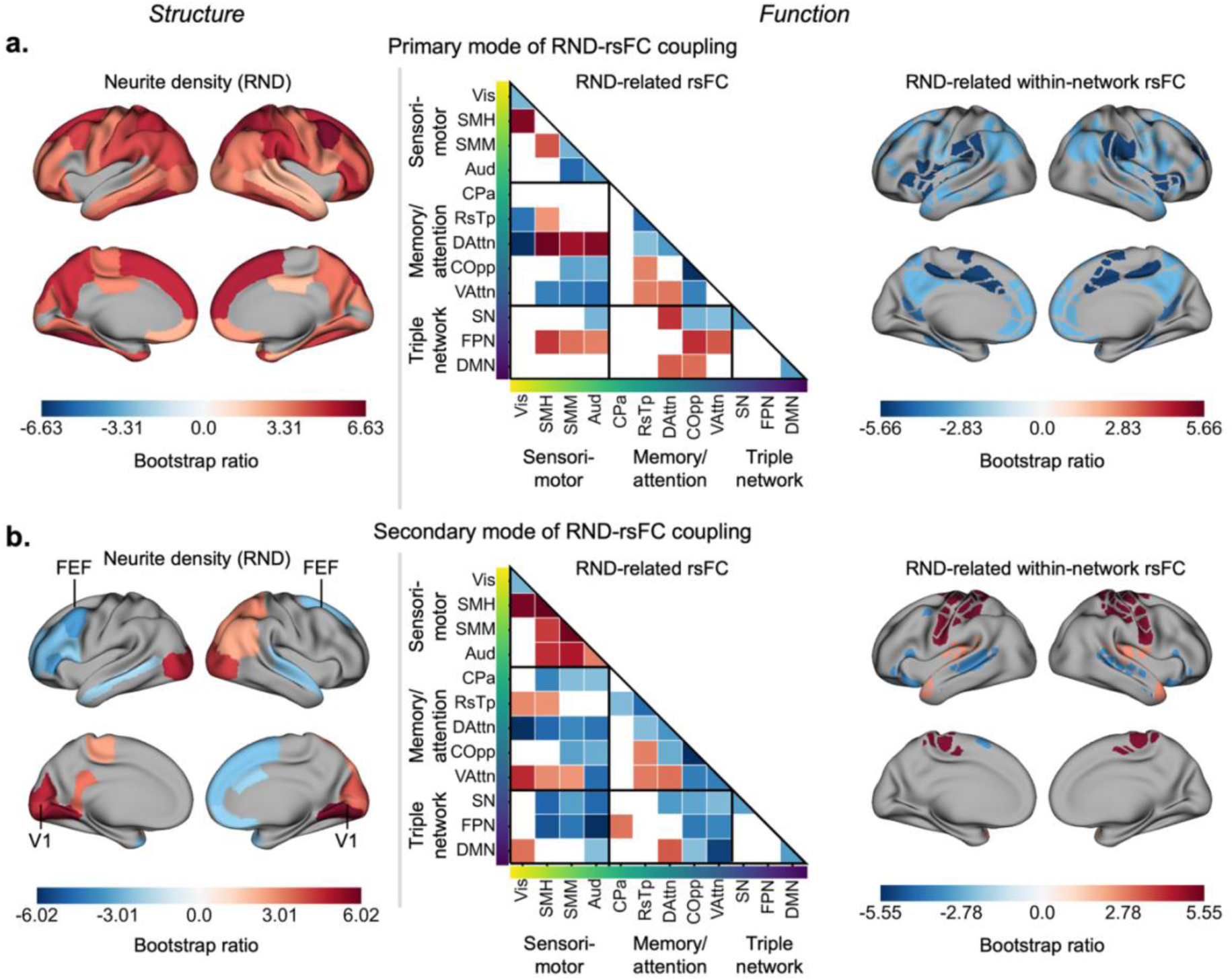
Primary and secondary patterns of coupling between neurite density and functional connectivity. Neurite density patterns are shown on the left, as bootstrap ratios representing brain regions that significantly contribute to the primary (a) and secondary (b) modes of RND-rsFC coupling. Related patterns of functional connectivity are shown in the middle (all rsFC) and right (within-network rsFC) as bootstrap ratios representing network connections that significantly contribute to the primary and secondary modes of RND-rsFC coupling. Networks are grouped based on their function into sensorimotor, attention/memory, and triple network model, which are ordered from unimodal to transmodal along the x-and y-axes in the middle column. A) The primary mode of RND-rsFC coupling accounted for 44% of the shared variance between cortical RND and functional network connectivity. A widespread pattern of higher neurite density (i.e., RNI) that included much of cortex–except for the insula, primary visual cortex, and portions of superior temporal and inferior frontal gyri–was related to a weaker within-network connectivity (right) and a mixture of stronger and weaker connectivity of dorsal and ventral attention networks, as well as frontoparietal network. B) The secondary mode of RNI-rsFC coupling accounted for 17% of the shared variance between cortical RND and functional network connectivity. A posterior-to-anterior pattern of higher cellular density in posterior regions (i.e., pericalcarine gyrus, V1; lateral occipital, right inferior parietal, left posterior cingulate) and higher cellular density in anterior regions (i.e., left rostral middle frontal gyrus; right superior frontal, orbitofrontal, and caudal anterior cingulate gyri) was related to a gradient of sensory and motor network connectivity that was largely stronger with other sensorimotor networks and weaker with the triple network mode, but mixed with memory/attention networks (e.g., weaker with cinguloparietal, dorsal attention; stronger with retrosplenial temporal, ventral attention). Abbreviations: visual, Vis; auditory, Aud; somatomotor-hand, SMH; somatomotor-mouth, SMM; dorsal attention, DAttn; ventral attention, VAttn; retrosplenial temporal, RsTp; cinguloparietal, CPa; cingulo-opercular, COpp; salience, SN; frontoparietal, FPN; default mode, DMN.

The secondary latent dimension of RND-rsFC coupling explained 17% of the shared variance between neurite density and resting-state connectivity, reflecting a posterior-to-anterior RND gradient with a sensorimotor rsFC gradient (Figure 3b). This pattern included greater posterior RND, i.e., in bilateral occipital, right parietal, and left paracentral and posterior cingulate gyri, but less anterior RND, i.e., in left superior temporal, medial superior frontal, mid-cingulate, and orbitofrontal gyri, as well as right middle and inferior frontal gyri (Figure 3b, left). This RND gradient was accompanied by a sensorimotor rsFC gradient (Figure 3b, middle), similar to the first latent dimension of RNI-rsFC coupling, that was characterized by stronger connectivity among sensorimotor networks, but weaker rsFC between sensorimotor and attention, memory, and higher-order networks, as well as weaker rsFC of higher-order networks in general. Likewise, RND-related within-network rsFC (Figure 3b, right) was stronger in sensorimotor networks (except visual) and weaker in attention and higher-order networks.

## Discussion

By bridging early cytoarchitectonic theory with modern multimodal neuroimaging approaches, this study identified patterns of cortical microarchitecture that support functional connectivity profiles in the preadolescent brain. Cellular density (i.e., RNI) coupling with functional connectivity was dominated by the primary pattern–whole-brain cellularity differences linked to a sensorimotor connectivity gradient. Conversely, neurite density (i.e., RND) coupling with functional connectivity lacked a single dominant pattern, instead exhibiting coexisting, regionally specific patterns. Two functional connectivity profiles were identified, differentially linked to these patterns of cellularity and neurite density. The first was a sensorimotor connectivity gradient that echoes prior research on functional brain development (Dong et al., 2021; Serio et al., 2026; Sydnor et al., 2023; Xia et al., 2022), which our findings link to global cellularity differences and a rostro-caudal neurite density gradient. The second centered on attention networks: stronger connectivity in top-down dorsal attention and frontoparietal networks contrasted with weaker connectivity in bottom-up ventral attention and cingulo-opercular networks, supported by the primary pattern of widespread greater neurite density and the secondary, rostro-caudal gradient of cellular density. Building on Brodmann’s hypothesis that cytoarchitectonic differences underlie functional specialization (Brodmann, 1909), these findings reveal how aspects of cortical microarchitecture differentially support preadolescent topology in sensorimotor and attentional networks.

### Cellular and neurite density patterns differ statistically and neuroanatomically

Combined *ex vivo* histological and *in vivo* MRI studies have found that cytoarchitectonically similar brain regions exhibit stronger connectivity than do less similar regions (Wei et al., 2018). Here, we estimated properties of cortical microarchitecture *in vivo* using RSI of diffusion-weighted MRI data, which is supported by histological validation (White et al., 2012). Restricted diffusion from RSI reflects intracellular spaces in brain tissue, with isotropic restricted diffusion (RNI) reflecting the presence of cell bodies (i.e., neuronal and glial; together, *cellulav density*) and directional restricted diffusion (RND) reflecting the presence of axons and dendrites (together, *neuvite density*) (White et al., 2012). Separate PLSC models reveal that the same amount of shared structure-function covariance is explained by a greater diversity of neurite density patterns compared to cellular density patterns. Essentially, most shared cellularity-rsFC variance is explained by a single latent pattern, but no single pattern of neurite-rsFC coupling accounts for more than 50% of shared variance. Instead, there are several coexisting patterns by which cortical neurite density is associated with functional connectivity differences. The primary direction of RND in cortex is radial with respect to the cortical surface, as are cortical columns (White et al., 2012), while genetic enrichment analyses link RND to genes involved in the developing and guiding neuronal projections (Fan et al., 2022), together suggesting that RND is a proxy for neurite density within cortical columns. Diverse patterns of local axon and dendrite architecture in preadolescents may facilitate regions’ specific long-range functional associations across the brain, compared with a single predominating pattern of cortical cell packing. One possible explanation for this distinction is that neuronal and glial cell bodies represent metabolic and structural needs that are more homogenous across cortex, while neurites represent hyperlocal connectional architecture that is specialized within functional units.

The primary and secondary patterns of rsFC-related cellular and neurite density were not identical but shared some common organization. For both, the primary patterns included most cortical regions, if not all, with the same directionality: greater cellular density across all cortical regions and greater neurite density in all cortical regions except pericalcarine, insular, left inferior frontal, and right paracentral gyri. The secondary patterns both exhibited rostro-caudal gradients, to some extent, that echo prior histological findings (Hilgetag et al., 2019). Cellular density was lower in caudal regions (e.g., V1, posterior cingulate, later occipital) but higher in rostral regions (e.g., anterior cingulate, insular, entorhinal cortices), while neurite density was the opposite (e.g., higher caudally in occipital, parietal gyri; lower rostrally in frontal gyri). Interestingly, the primary cellular density pattern and secondary neurite density pattern were both linked to a connectivity gradient in sensorimotor networks, while the primary neurite density pattern and secondary cellular density pattern were both linked to similar attention network connectivity. If RND-estimated neurite density represents the density of hyperlocal connectivity (i.e., of axons and dendrites within cortical columns), this suggests that the sensorimotor-association gradient that describes developmental timing (Gogtay et al., 2004; Sydnor et al., 2023) and adult functional connectivity (Margulies et al., 2016) is partially underscored by rostro-caudal differences in local synaptic architecture.

### Common network connectivity profiles between neurite and cellular density

Finally, we found two notably similar network connectivity profiles linked to both cellular and neurite density.

First, a sensorimotor connectivity gradient was characterized by stronger connectivity among sensory and motor networks (i.e., visual, auditory, somatomotor-hand and -mouth) and weaker connectivity between sensorimotor and higher-order networks in the triple network model (i.e., salience, frontoparietal, default mode). Prior studies of connectivity gradients have uncovered a principal gradient of network organization with sensory and motor regions at one pole and higher-order, associative networks (e.g., limbic, default mode) at the other (Huntenburg et al., 2018; Margulies et al., 2016; Sydnor et al., 2021). More recently, longitudinal studies of child brain network development suggest that this gradient emerges during the transition to adolescence (Dong et al., 2021), in part due to pubertal maturation (Gracia-Tabuenca et al., 2021; Serio et al., 2026). The sensorimotor connectivity gradient identified here may represent the immature state of this gradient in preadolescence, in which early developing sensory and motor networks exhibit a gradient of connectivity that differs between sensorimotor networks and higher-order associative networks in the triple network model, while these later-developing associative higher-order networks do not yet exhibit varying connectivity along this axis. Interestingly, this sensorimotor gradient was linked to the dominant, primary pattern of greater cellular density across cortex, but to the secondary, rostro-caudal pattern of neurite density: lower in posterior regions (e.g., primary visual, parietal, posterior cingulate) and higher in anterior regions (e.g., frontal eye fields, superior frontal gyrus, anterior cingulate). Together with the considerable difference in covariance explained by the primary cellular and secondary neurite density patterns, this distinction may suggest that, in the developing brain, the emerging dominant gradient of functional network organization is largely supported by differences in cellular density across the cortex and, to a lesser extent, by a rostro-caudal pattern of neurite density differences. That is, the density of cell bodies (e.g., neuronal cell bodies, glia) may be more relevant for this major gradient of long-range cortical communication than the density or organization of cortical neurites (e.g., axons and dendrites in cortical columns).

Second, an attention network profile was characterized by contrasting connectivity between different aspects of attentional control (Dosenbach et al., 2025; Seeley et al., 2007; Vossel et al., 2014). The top-down, goal-directed dorsal attention and frontoparietal networks exhibited stronger connectivity, while bottom-up, action-oriented ventral attention and cingulo-opercular networks exhibited weaker connectivity. This contrast aligns with developmental trends that emerge during adolescence, as stimulus-driven distraction is tempered by improving executive control (B. Casey et al., 2016; Kang et al., 2022; Vetter et al., 2015). Attentional and control systems may serve as important regulators of functional connectome maturation throughout adolescence (Dong et al., 2024; Keller et al., 2023) and disruption or imbalance in these systems poses transdiagnostic risk for psychopathology (Dziemian et al., 2025; Eysenck et al., 2007; Li et al., 2026; Morea & Calvete, 2021; Shadur & Lejuez, 2015). Notably, our findings suggest that this important preadolescent connectivity profile is supported by widespread cortical neurite density and, to a lesser extent, by a rostro-caudal pattern of cellular density. Thus, the hyperlocal (i.e., intraregional) connectional architecture represented by neurite density may support the maturation of functional brain organization and play a role in the adolescent emergence of psychopathology.

These connectivity profiles and their differential support from widespread primary and rostro-caudal secondary microarchitecture patterns suggest a reciprocal mapping between cortical microarchitecture and functional network organization. While macroscale sensorimotor communication is anchored primarily by widespread cortical cell packing, attention networks depend more on regional axonal and dendritic density, supported only secondarily by regional differences in cell body density. Future, longitudinal research should follow these patterns across adolescence to test developmental stability of this structure-function coupling and determine the role of later-developing networks, including the triple network model. Ultimately, investigating individual differences in these microarchitecture and functional connectivity patterns may provide additional insight into their role in healthy socioemotional maturation and the adolescent emergence of psychopathology.

### Strengths & Limitations

This study capitalizes on the strengths of a large, rich neuroimaging dataset to identify cross-sectional patterns of microarchitecture-function coupling in preadolescents but also faces a few limitations. The sample analyzed here includes over 6,000 youth ages 9-10 years from across the United States at a key preadolescent inflection point where pubertal influences are beginning to influence brain development. Future, longitudinal work with multiple, large-scale neuroimaging datasets should investigate how the identified patterns of structure-function coupling change across puberty and whether these patterns are universal, across sociodemographic and geopolitical boundaries. RSI provides histologically validated estimates of gray matter microarchitecture and using a cortical brain atlas provides anatomically driven dimensionality reduction that likely increases the interpretability of identified coupling patterns. However, the regional estimates of cortical microarchitecture used here likely span multiple cytoarchitectonic areas (Amunts & Zilles, 2015; Devlin & Poldrack, 2007). Thus, the “expression” of coupling patterns identified here may reflect anatomical differences in cytoarchitectonic boundaries as much as they reflect microarchitecture differences within regions. Individual-level, multi-modal data fusion could help distinguish these two potential sources of individual differences in microarchitecture-function coupling patterns and should be explored in future work (Sui et al., 2023). Finally, it was beyond the scope of this work to assess how individual differences in microarchitecture-function coupling are related to other aspects of development (e.g., puberty), behavior, cognition, or social and environmental factors. Future work should further investigate the coupling patterns identified here to better understand their role in child and adolescent development.

## Conclusions

Using multimodal neuroimaging in a large, nationwide cohort of over 6,000 9–10-year-old children, this study identified differential microarchitecture patterns underlying sensorimotor and attentional functional connectivity profiles in the preadolescent brain. Specifically, we identified a microstructural dissociation by which a sensorimotor connectivity gradient is supported by one dominant, global pattern of cellular density and a profile of contrasting attention network connectivity is supported by widespread neurite density, though both connectivity profiles are secondarily supported by rostro-caudal gradients of neurite and cellular density, respectively. Identifying these modes of structure-function coupling in the preadolescent brain, at the precipice of pubertal development and adolescent brain maturation, provides a foundation for future longitudinal study of individual differences in these microarchitecture patterns and functional connectivity profiles and how they explain adolescent cognitive development and the emergence of psychopathology.

## Acknowledgments

A special thank you to all of the children and families for their participation in their ABCD Study.

Research described in this article was supported by the National Institutes of Health (MMH: NIEHS R01ES032295, R01ES031074; KLB: R00MH135075; JM: F31HD122332) CCI would like to acknowledge scholars involved in NSP (R25 NS089462), BRAINS (R25 NS094094), and Diversifying CNS (R25 NS117356), as well as R25MH125545 and R25MH120869 for creating a supportive network of ABCD Study users.

Data used in the preparation of this article were obtained from the Adolescent Brain Cognitive Development^SM^ (ABCD) Study (https://abcdstudy.org), held in the NIMH Data Archive (NDA). This is a multisite, longitudinal study designed to recruit more than 10,000 children age 9-10 and follow them over 10 years into early adulthood. The ABCD Study® is supported by the National Institutes of Health and additional federal partners under award numbers U01DA041048, U01DA050989, U01DA051016, U01DA041022, U01DA051018, U01DA051037, U01DA050987, U01DA041174, U01DA041106, U01DA041117, U01DA041028, U01DA041134, U01DA050988, U01DA051039, U01DA041156, U01DA041025, U01DA041120, U01DA051038, U01DA041148, U01DA041093, U01DA041089, U24DA041123, U24DA041147. A full list of supporters is available at https://abcdstudy.org/federal-partners.html. A listing of participating sites and a complete listing of the study investigators can be found at https://abcdstudy.org/consortium_members/. ABCD consortium investigators designed and implemented the study and/or provided data but did not necessarily participate in the analysis or writing of this report. This manuscript reflects the views of the authors and may not reflect the opinions or views of the NIH or ABCD consortium investigators. The ABCD data repository grows and changes over time. The ABCD data used in this report came from https://dx.doi.org/10.15154/z563-zd24.

## Author Contributions

Conceptualization: KLB, MMH

Data curation: KLB, KS

Formal Analysis: KLB, KS

Funding acquisition: MMH, KLB

Methodology: KLB, MMH

Project administration: KLB, MMH

Resources: MMH

Software: KLB, KS

Supervision: MMH

Visualization: KLB

Writing – original draft: KLB, MMH, JM

Writing – review & editing: KLB, KS, MMH, JM, CCI

