## Supplementary Material for "Cortical microarchitecture supports preadolescent functional brain network connectivity"

### Local cortical microarchitecture underlying macroscale functional connectivity in late childhood

This document contains:

**Supplementary Methods** **3**

Supplementary Figure 1. Participant inclusion and exclusion flowchart. 3

**Supplementary Results** **4**

Additional RNI-rsFC latent dimensions 4

Supplementary Figure 2. Third pattern of RNI-rsFC coupling (3% of RNI-rsFC covariance). 4

Supplementary Figure 3. Fourth pattern of RNI-rsFC coupling (3% of RNI-rsFC covariance). 4

Supplementary Figure 4. Fifth pattern of RNI-rsFC coupling (2% of RNI-rsFC covariance). 5

Supplementary Figure 5. Sixth pattern of RNI-rsFC coupling (1% of RNI-rsFC covariance). 5

Supplementary Figure 6. Seventh pattern of RNI-rsFC coupling (1% of RNI-rsFC covariance). 6

Supplementary Figure 7. Eighth pattern of RNI-rsFC coupling (1% of RNI-rsFC covariance). 6

Supplementary Figure 8. Ninth pattern of RNI-rsFC coupling (1% of RNI-rsFC covariance). 7

Supplementary Figure 9. Tenth pattern of RNI-rsFC coupling (1% of RNI-rsFC covariance). 7

Additional RND-rsFC latent dimensions 7

Supplementary Figure 10. Third pattern of RND-rsFC coupling (6% of RND-rsFC covariance). 7

Supplementary Figure 11. Fourth pattern of RND-rsFC coupling (6% of RND-rsFC covariance). 8

Supplementary Figure 12. Fifth pattern of RND-rsFC coupling (4% of RND-rsFC covariance). 8

Supplementary Figure 13. Sixth pattern of RND-rsFC coupling (4% of RND-rsFC covariance). 9

Supplementary Figure 14. Seventh pattern of RND-rsFC coupling (3% of RND-rsFC covariance). 9

Supplementary Figure 15. Eighth pattern of RND-rsFC coupling (3% of RND-rsFC covariance). 10

Supplementary Figure 16. Ninth pattern of RND-rsFC coupling (2% of RND-rsFC covariance). 10

*Corresponding Author:* Dr. Katherine L. Bottenhorn, Assistant Professor of Psychology, Florida International University, Miami, FL, USA


### Supplementary Methods

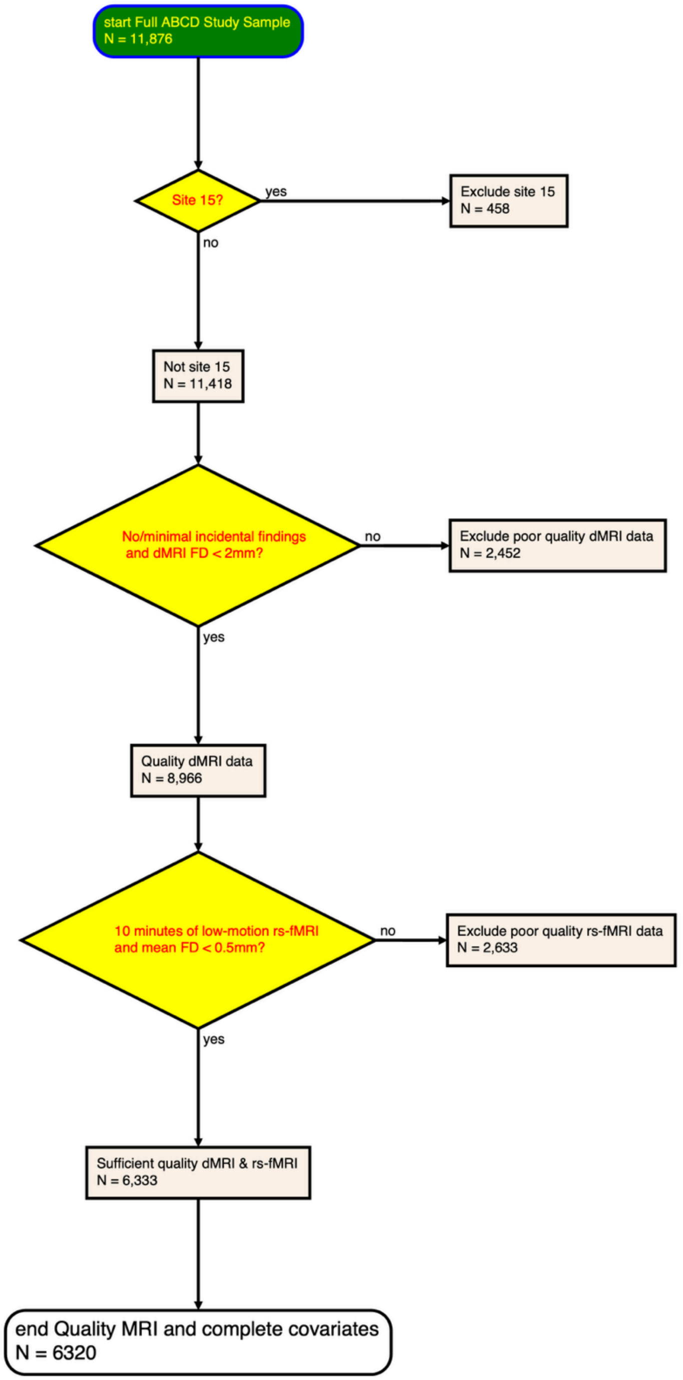


***Supplementary Figure 1. Participant inclusion and exclusion flowchart.***
Specifies the number of participants (*N*) included and excluded at each stage. Exclusion criteria for the ABCD study included lack of English proficiency, severe sensory, neurological, medical or intellectual limitations, and inability to complete an MRI scan. For this study, we first excluded individuals from site 15 due to hardware issues (*N* = 458); then further

excluded participants who had incidental neurological findings from their MRI scans, failed the ABCD Study imaging quality control procedures, or had greater than 2mm of head motion during dMRI scans ($N$ = 2,452); and finally excluded participants who had greater than 0.5mm head motion during rs-fMRI scans ($N$ = 2,633) for a final sample of 6,320 participants with sufficient quality dMRI and rs-fMRI data and complete data on all covariates. The final sample characteristics for the current study are described in Table 1.

### Supplementary Results

For the following figures, cortical microarchitecture (i.e., cellular density, RNI, or neurite density, RND) patterns are shown on the left, as bootstrap ratios representing brain regions that significantly contribute to latent dimensions, or modes, of structure-function coupling. Significant, related patterns of functional connectivity are shown in the middle column (i.e., as an adjacency matrix representing all within- and between-network rsFC) and right column (i.e., within-network rsFC only, on brain surfaces to show topography) as bootstrap ratios representing network connections that significantly contribute to latent dimensions, or modes, of structure-function coupling. Networks in the middle column's adjacency matrices are ordered along the sensorimotor-association axis as in the main text.

Abbreviations for the additional latent dimension figures below are as follows:

- RNI: restricted normalized isotropic diffusion, spherical intracellular diffusion reflecting cellular density
- rsFC: resting-state functional connectivity
    - Vis: visual network
    - SMH: somatomotor-hand network
    - SMM: somatomotor-mouth network
    - Aud: auditory network
    - CPa: cinguloparietal network
    - RsTp: retrosplenial temporal network
    - DAttn: dorsal attention network
    - COpp: cingulo-opercular network
    - VAttn: ventral attention network
    - SN: salience network
    - FPN: frontoparietal network
    - DMN: default mode network

#### Additional RNI-rsFC latent dimensions

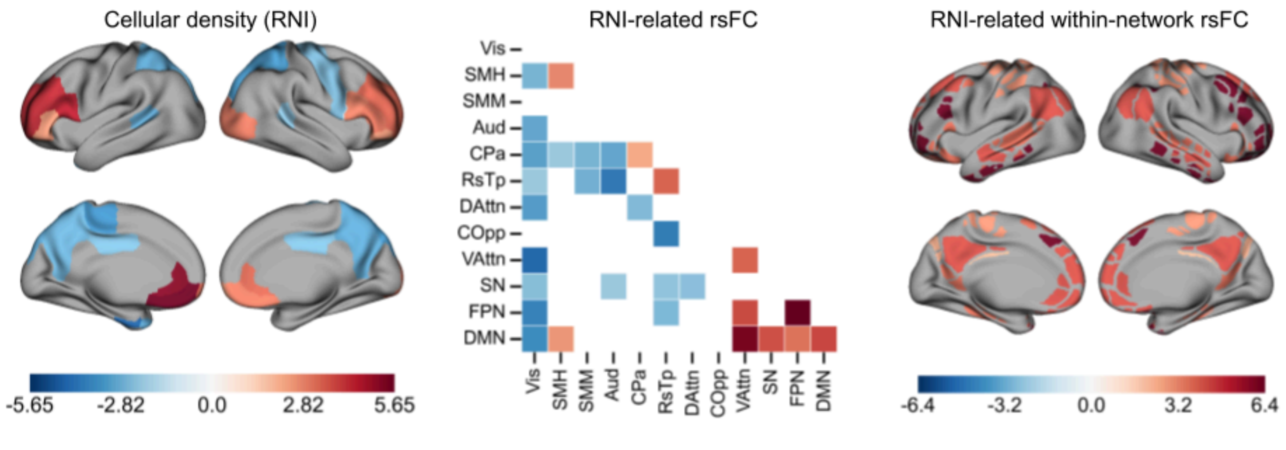


***Supplementary Figure 2. Third pattern of RNI-rsFC coupling (3% of RNI-rsFC covariance).***

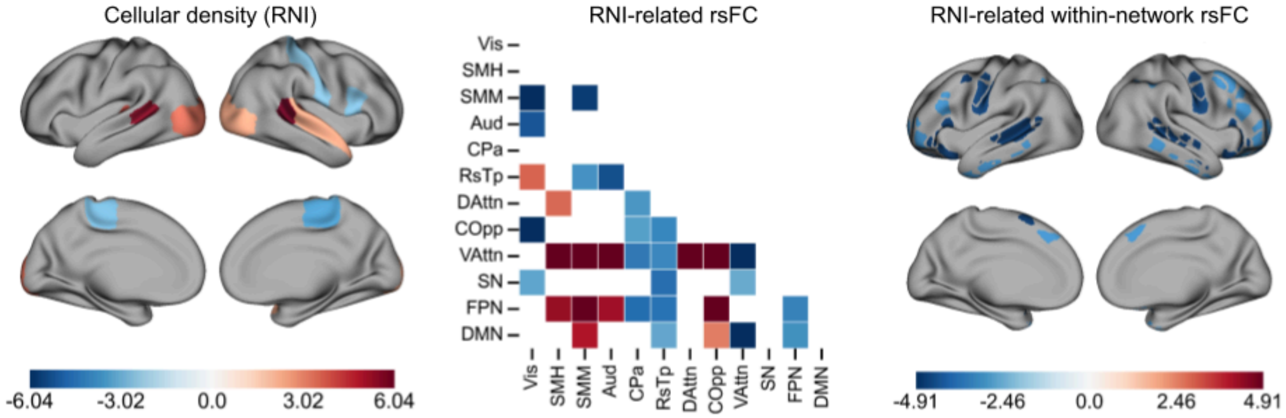


***Supplementary Figure 3. Fourth pattern of RNI-rsFC coupling (3% of RNI-rsFC covariance).***

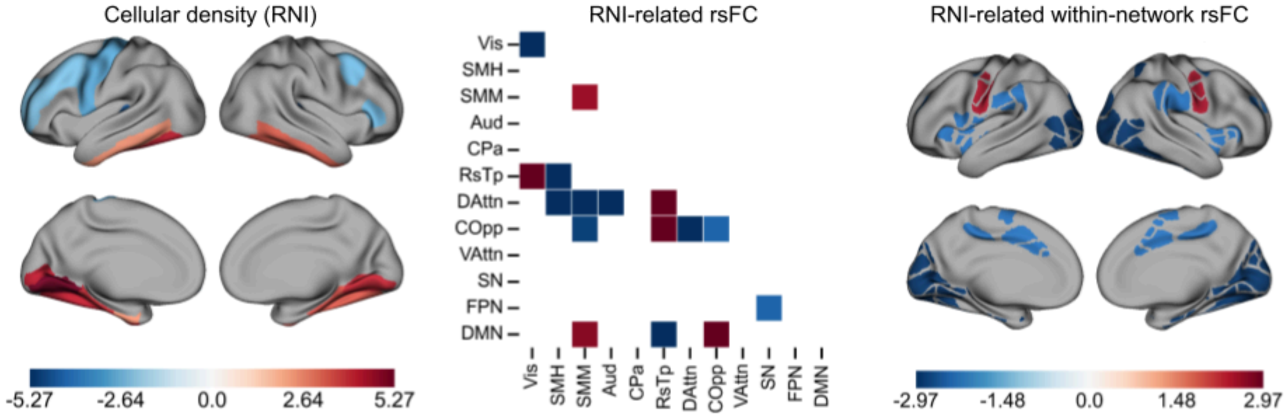


***Supplementary Figure 4. Fifth pattern of RNI-rsFC coupling (2% of RNI-rsFC covariance).***

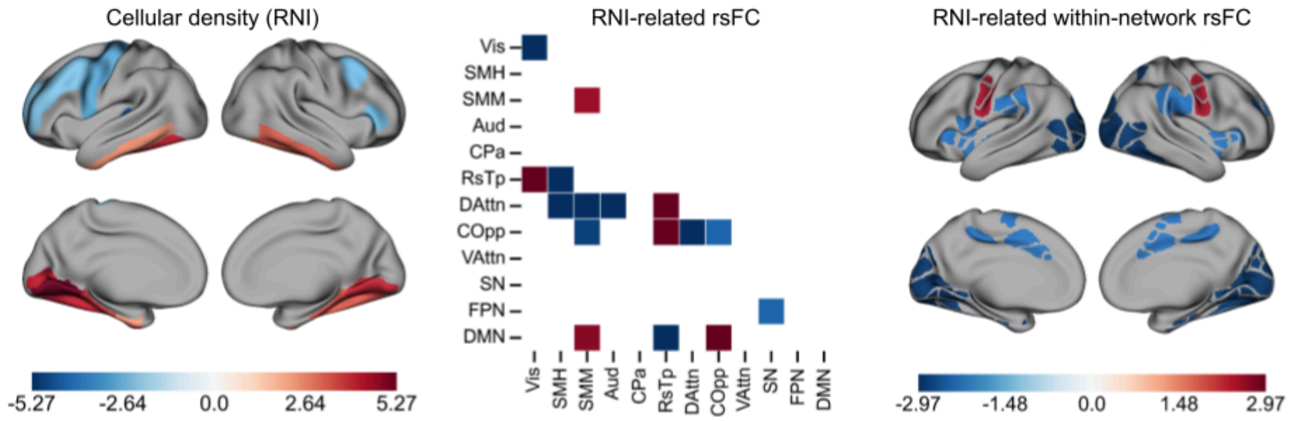


***Supplementary Figure 5. Sixth pattern of RNI-rsFC coupling (1% of RNI-rsFC covariance).***

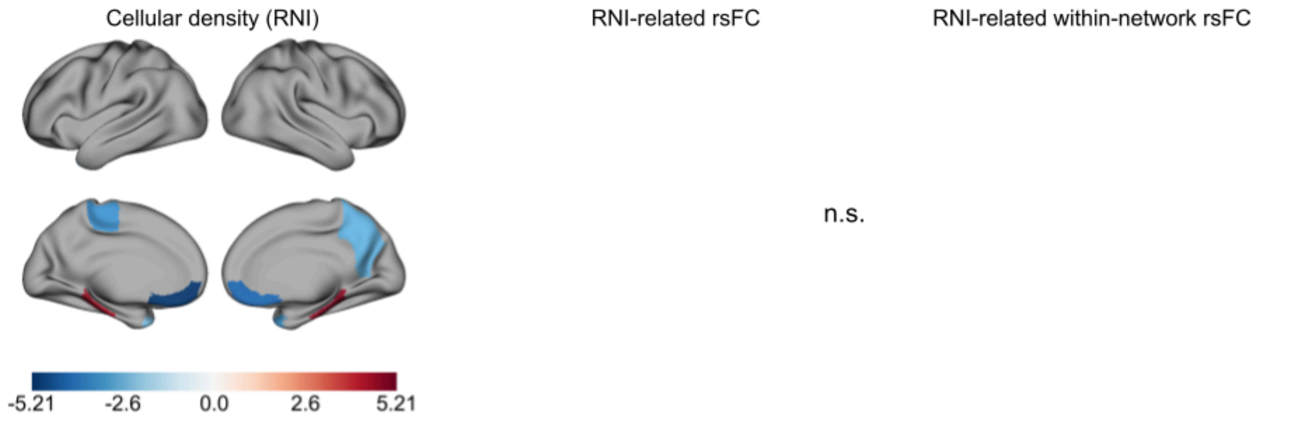


***Supplementary Figure 6. Seventh pattern of RNI-rsFC coupling (1% of RNI-rsFC covariance).***
A lack of significant RNI-related rsFC suggests that this pattern of cellular density may be diffusely related to rsFC, but that no specific connections are driving this pattern.

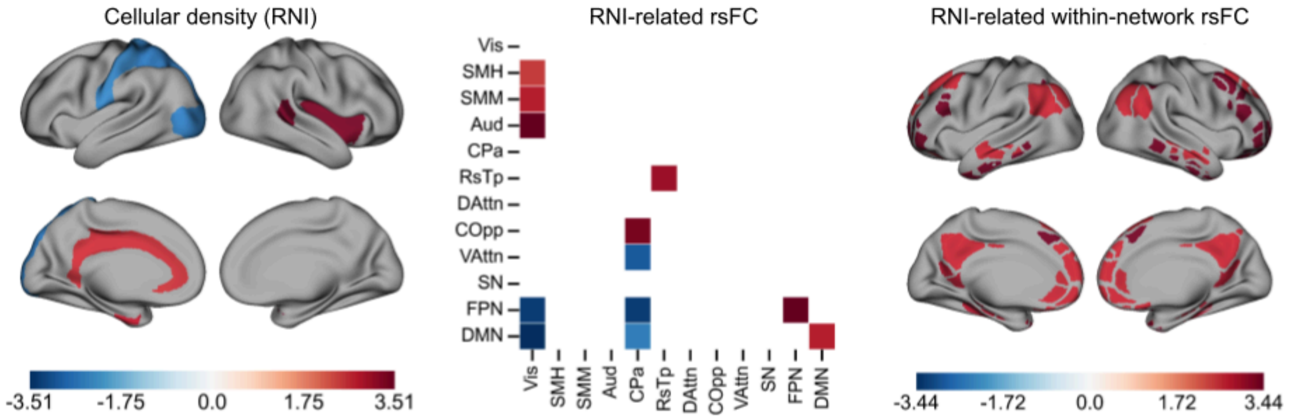


***Supplementary Figure 7. Eighth pattern of RNI-rsFC coupling (1% of RNI-rsFC covariance).***

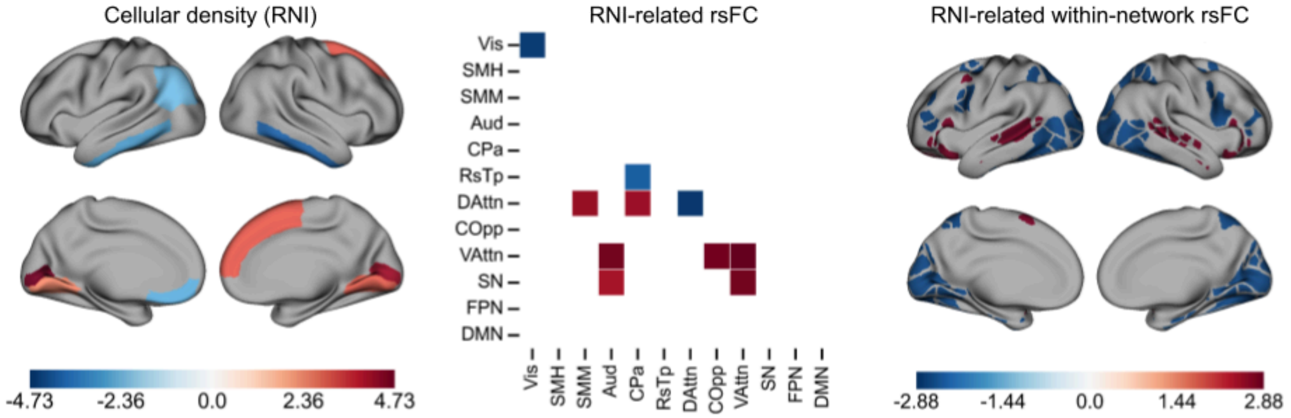


***Supplementary Figure 8. Ninth pattern of RNI-rsFC coupling (1% of RNI-rsFC covariance).***

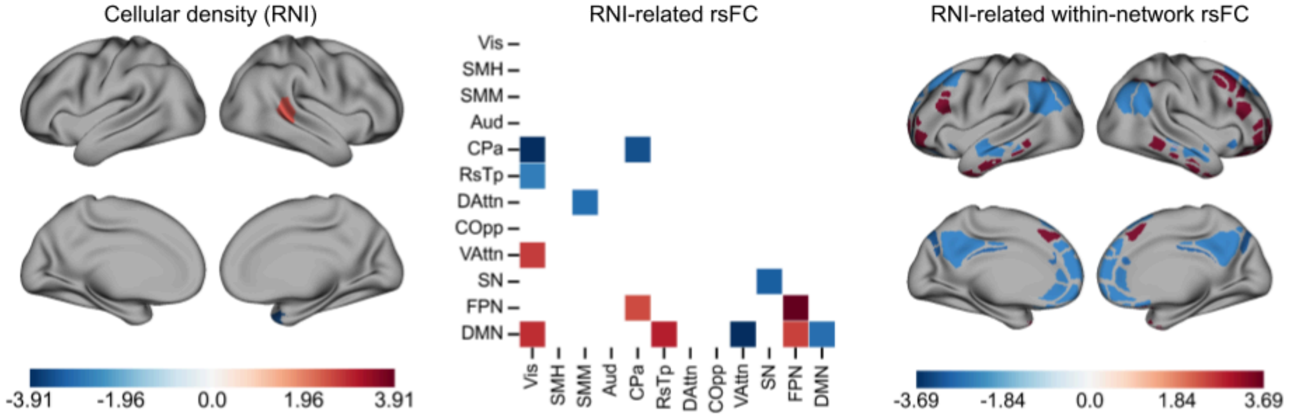


***Supplementary Figure 9. Tenth pattern of RNI-rsFC coupling (1% of RNI-rsFC covariance).***

#### Additional RND-rsFC latent dimensions

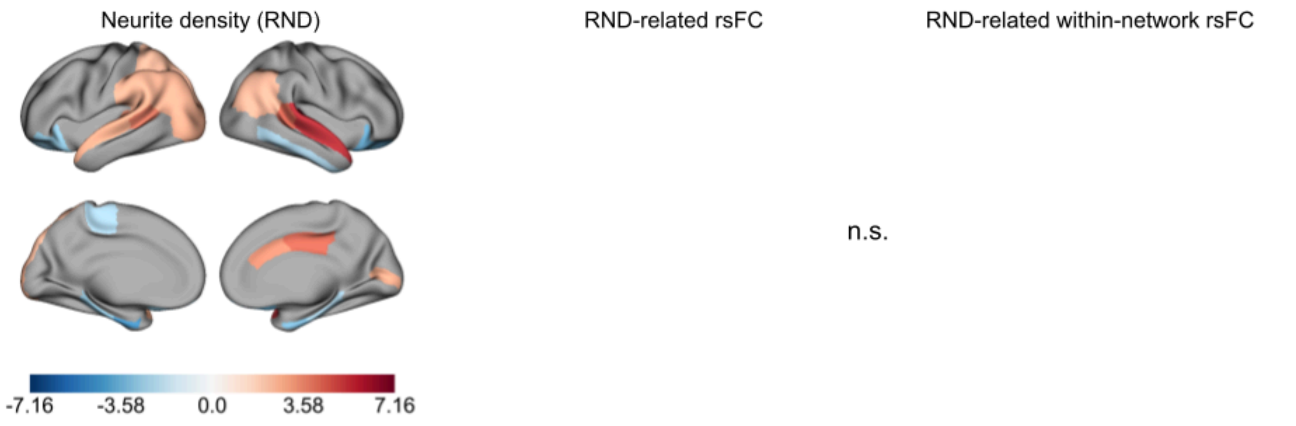


***Supplementary Figure 10. Third pattern of RND-rsFC coupling (6% of RND-rsFC covariance).***
A lack of significant RND-related rsFC suggests that this pattern of neurite density may be diffusely related to rsFC, but that no specific connections are driving this pattern.

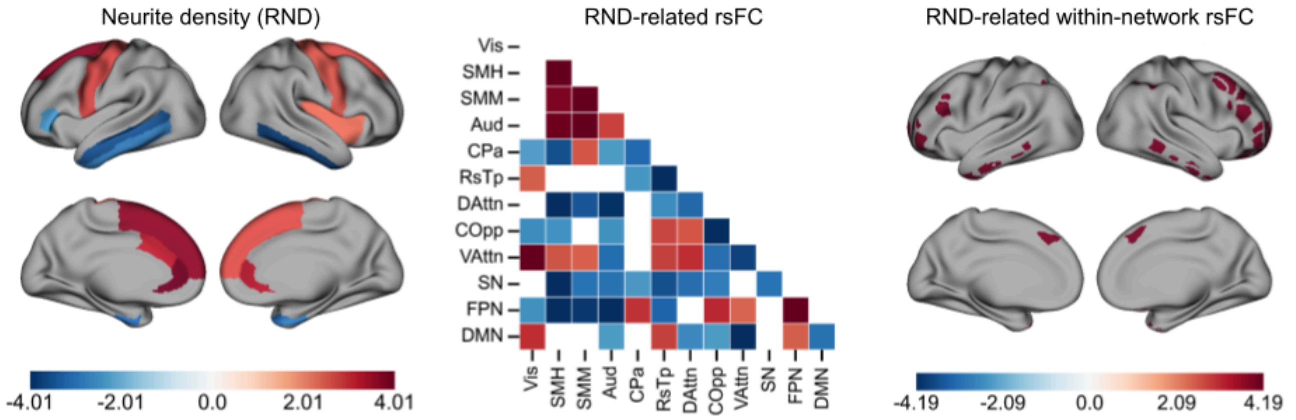


***Supplementary Figure 11. Fourth pattern of RND-rsFC coupling (6% of RND-rsFC covariance).***

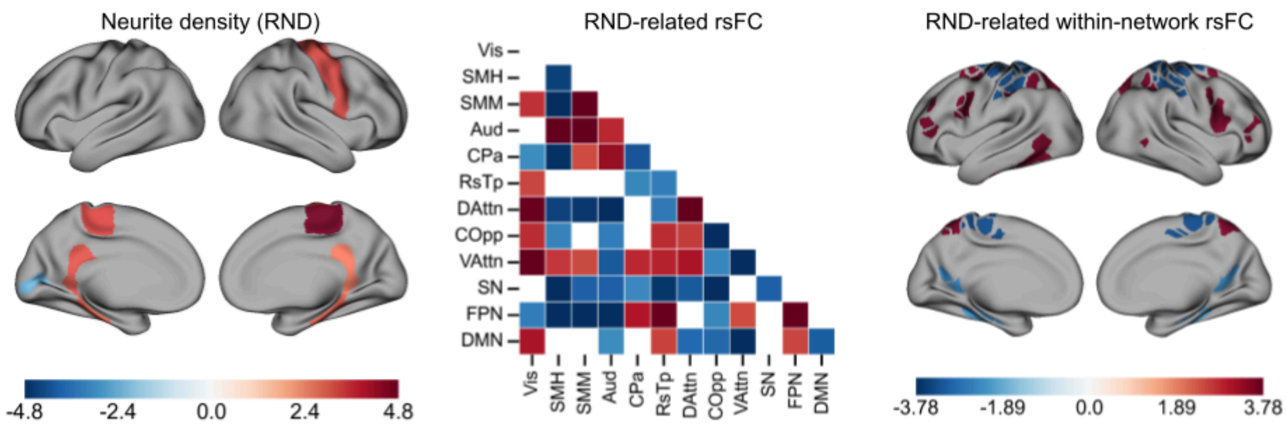


*Supplementary Figure 12. Fifth pattern of RND-rsFC coupling (4% of RND-rsFC covariance).*

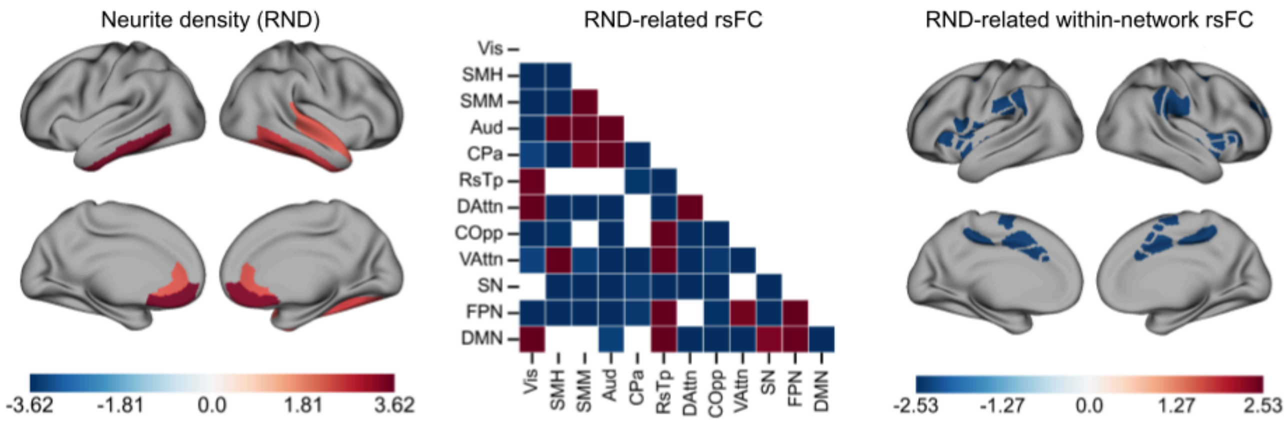


*Supplementary Figure 13. Sixth pattern of RND-rsFC coupling (4% of RND-rsFC covariance).*

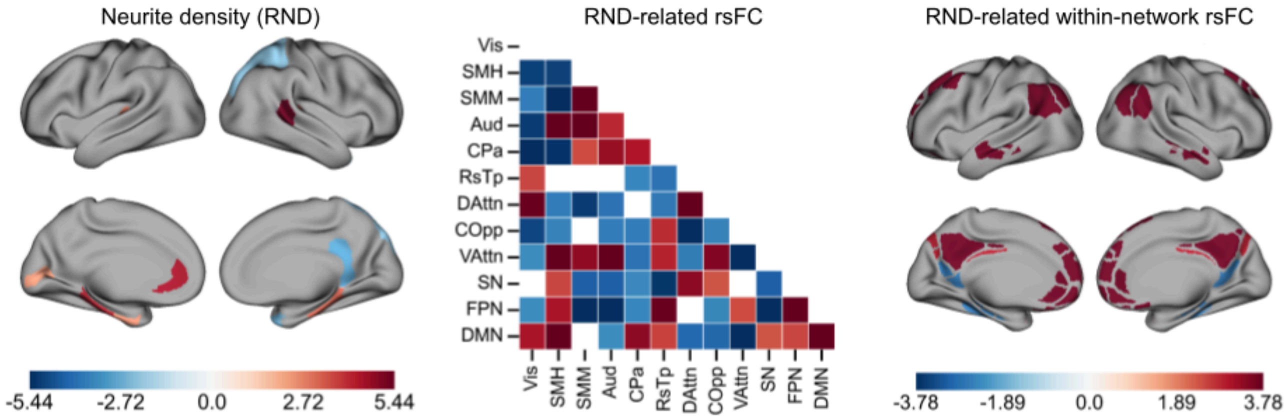


***Supplementary Figure 14. Seventh pattern of RND-rsFC coupling (3% of RND-rsFC covariance).***

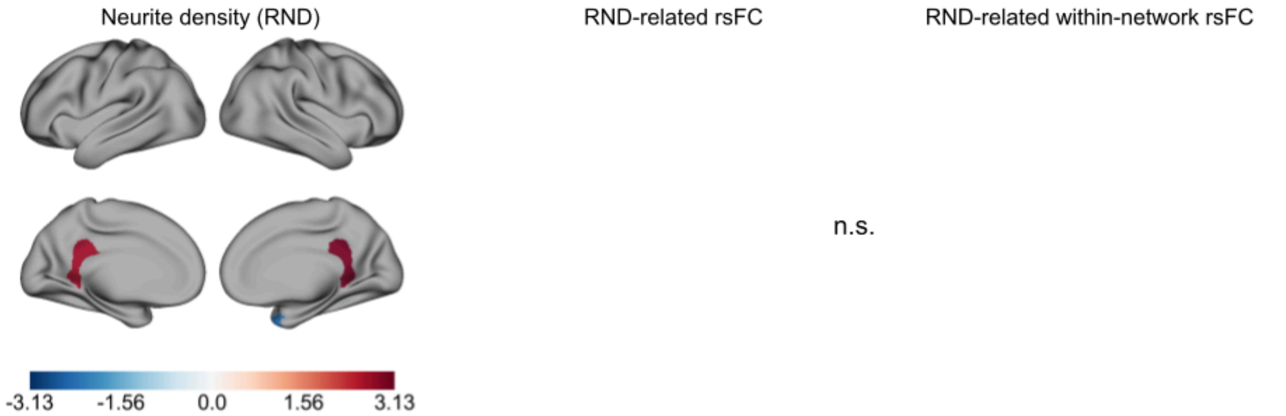


***Supplementary Figure 15. Eighth pattern of RND-rsFC coupling (3% of RND-rsFC covariance).***
A lack of significant RND-related rsFC suggests that this pattern of neurite density may be diffusely related to rsFC, but that no specific connections are driving this pattern.

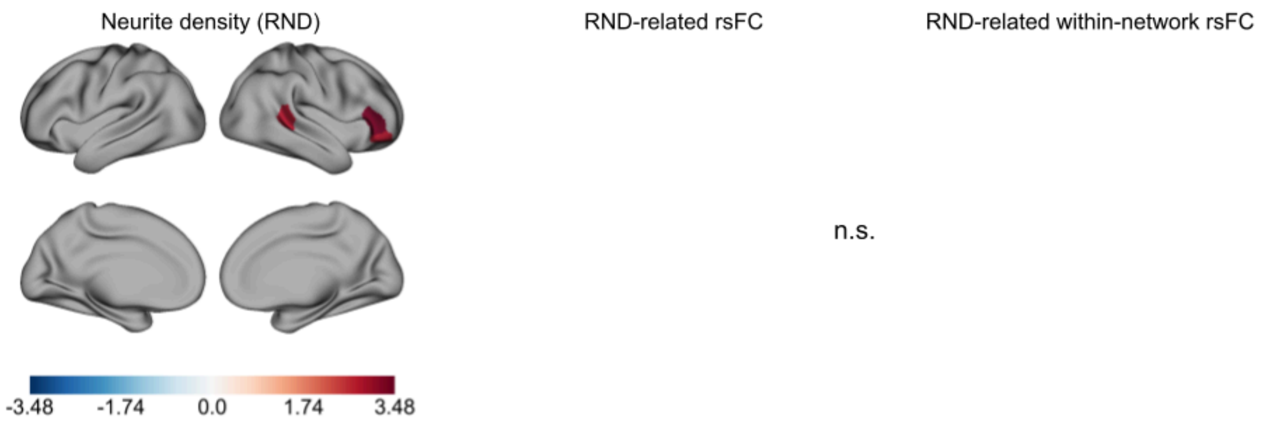


***Supplementary Figure 16. Ninth pattern of RND-rsFC coupling (2% of RND-rsFC covariance).***
A lack of significant RND-related rsFC suggests that this pattern of neurite density may be diffusely related to rsFC, but that no specific connections are driving this pattern.
